# c-Fos protein shRNA blockade in the central amygdala nucleus interfere with rats emotional reactivity on behavioral and autonomic level

**DOI:** 10.1101/2022.01.18.476659

**Authors:** Ita Robakiewicz

## Abstract

This report is focusing on a function of the c-Fos protein in an associative, stress-induced memory. The shRNA vector injections were utilised to functionally silence the central amygdala nucleus in adult Wistar rats. Subsequently the operated animals and their control counterparts were screened in a selection of an emotionally-dependent tests and in a few standard behavioral neuroscience tools. Rats from the c-Fos silencing (ShFos) group expressed contra-depressive-like behaviors in Porsolt Swimming Test, spending more time actively searching for escape way then the rats from the control group. ShFos rats also had engaged in a more rapid activity in the Open Field Test, showing a decline in the neo-phobia. Micturition was decreased in shFos animals, indicating a change in the emotionality on an autonomic level. Presented results are showcasing a multi-directional regulation of the behaviors from the central amygdala nucleus by the c-Fos activity.

## Introduction

The ensemble of subcortical structures jointly named amygdala (Amy), is known to be an emotional hub of the brain and considered determinant of matching particular and current experiences with the emotional valence (Calhoon and Tye, 2015), that also applies to the resurfacing memories as well (Ciocchi et al., 2010). The complex functions of the Amy are characterised by its wide projections. Two main Amy nuclei are: (1) the basolateral nucleus (BLA) being composed mainly of the glutamate, excitatory neurons that are connecting to the cortical areas, as well as to the hypothalamus and to the limbic structures (hippocampus, nucleus accumbens) and (2) the central nucleus (CeA) connected mainly to the BLA as an inhibitory GABA-based controller and to the cortical areas, as well as the ventral hippocampus, periaqueductal gray (PAG) and the basal nuclei (Beyer and Dobrowska, 2020).

Changed activation of Amy as whole in human studies (Roozendaal et al., 2009) and its nuclei in particular in both human and animal studies (Fox et al., 2015) is symptomatic for psychological and behavioral maladaptive disorders. Changes are observed in affective disorders (Duvarci and Pare, 2014), spectrum of anxiety disorders (Babaev et al., 2018) and neurodegenerative disorders (Tzu-Wei et al., 2015).

Knowing that Amy and its functional connectivity are highly adaptively important, it is crucial to understand Amy’s nature from the molecular to the behavioral level.

Here the CeA nucleus is of an exceptional interest, since it was previously perceived as highly homogenous (Ehrlich et al., 2009), but the contemporary results prove otherwise – the CeA has an important role in generating behavioral output on the autonomic and complex levels. The subpopulations of neurons in the CeA, were described for their involvement in the detailed emotional gradient accretion (Nieh et al., 2013) in certain behavioral conditions for example: an inhibitory avoidance memory consolidation (Ghiasvand et al. 2011), generalised anxiety (Yan-Mei et al., 2019, Meyza et al., 2009), control of an aversive memory strength (Ozawa et al., 2016), stress-induced anxiety (Ventura-Silva et al., 2020), behavioral response to a challenge (Regev et al., 2012), conditioned fear (Wiktorowska et al. 2021, Campese et al., 2015), active avoidance (Martinez et al., 2013) and momentary arrests during exploration (Botta et al., 2020).

CeA activity is context dependent – distinct neuron populations can gate for approaching or avoidance behavior, based on external information (Ciocchi et al, 2010, Adraka et al., 2021, Warolw et al., 2017 and 2020, Lebitko et al., 2021). If one aims at changing the function of Amy (or its substructures) by different means (e.g. chemogenic, optogenetic, magnetic, electrical or mechanistic) they will need to tailor that means, to a specific characteristics of the receiver unit, in order to obtain effects on the behavioral level (Badrinarayan et al., 2012, Warolw et al., 2017 and 2020).

The c-Fos labelling in the brain specimens is regarded as activity marker, especially sensitive to novel stimulus or a novelty elements in the former known situations and set-ups. It was frequently and consistently described as Amy emotionally-dependent processing marker (Knapska et al., 2006 and 2007, Cho et al., 2017, Choi et al., 2013, De Francesco et al., 2015, Lam et al., 2018, Morland et al., 2016, Yanagida et al., 2016). It was proven that there is a c-Fos activation in CeA nucleus in the forced swimming task (Choi et al., 2013, De Francesco et al., 2015, Lam et al. 2018, Sherwin et al., 2017, Yanagida et al., 2016).

In this set of experiments a shRNA vector was used to functionally knockdown the CeA in rats, as for evaluation of the structure’s behavioral gating in Porsolt Swimming Test (PST). There were no former attempts of doing the latter. In order to characterise the locomortor, cognitive and the emotional functioning of the operated rats a standard set of behavioral neuroscience tools and a designed in our laboratory self-exposure paradigm have been used (Robakiewicz et al., 2019).

## Methods

### Subjects

21 male Wistar rats, 3 m.o.

### Stereotactic operations

The stereotactic operations were conducted on rats in respective manner: each animal was pre-anesthetized in hermetic box with isofluran (2 l/ min; Baxter; 5%). After the animal was showing no reflexes, it was put in the stereotactic frame, attached to the isofluran anestesia apparatus (2% and 0,4l/min). Analgesic butomidor was injected subcutaneously (2 mg/kg) in order to deepen the anaesthesia, before drilling the skull. Eyes of each animal were covered with petrolatum in order to prevent drying of the eyeballs. Scalp was shaved and sterilised with 70% ethyl alcohol, then dissected, cleaned of periosteum and visible vessels. Scalp was exposed and bregma and lambda were localised. Due to bregma and the stereotactic atlas of rat brain coordinates (Paxions & Watson, 2007), drilling placements were identified. In each hemisphere there was one hole drilled, 0.2 cm in anterior manner from bregma and +/- 0.38 cm in medio-lateral axis from bregma, in order to penetrate the brain and reach the CeA. Microinjections in 0.81 cm in depth from the skull surface, were carried out with Nanofil 10 μL syringe and UMP-3 controller attached to it (World Precision Instruments, Inc.).

Each animal was injected in each hole with 500 nL vector solution (in Phosphate-buffered saline - PBS) containing of ^~^0,6 x 10^9 of the virus particles with experimental construct silencing c-Fos (LV-GFP-sh-c-Fos) or the control construct silencing luciferase (LV-GFP-Sh-Luc). After the injections the needle maintained in the brain for 15 min latency to let the virus particles diffuse in the tissue as much as possible. After that period needle was removed and after the second injection, the scalp was sawed with surgical thread. Next the rats were awaken and moved to the home cages for convalescence period. The production process and the effectiveness of the viral constructs that were used in this study, was previously described by other researchers (De Hoz et al., 2018, Lebitko et al., 2021). The vectors were produced in the Laboratory of Animal Models in the Nencki Institute of Experimental Biology, Polish Academy of Sciences.

The accuracy of the injection sites was confirmed by the microscopic inspection of the presence of the green fluorescence marker (GFP), that was tagging the transfected neurons in the brain slices collected from all the animals (cut on cryostat-45 μm tick coronal sections, previously fixed in 4% paraformaldehyde in PBS and cryoprotected in 30% sucrose). The graphic description of the injection sites is in figure 1B and C.

**Figure 1.**
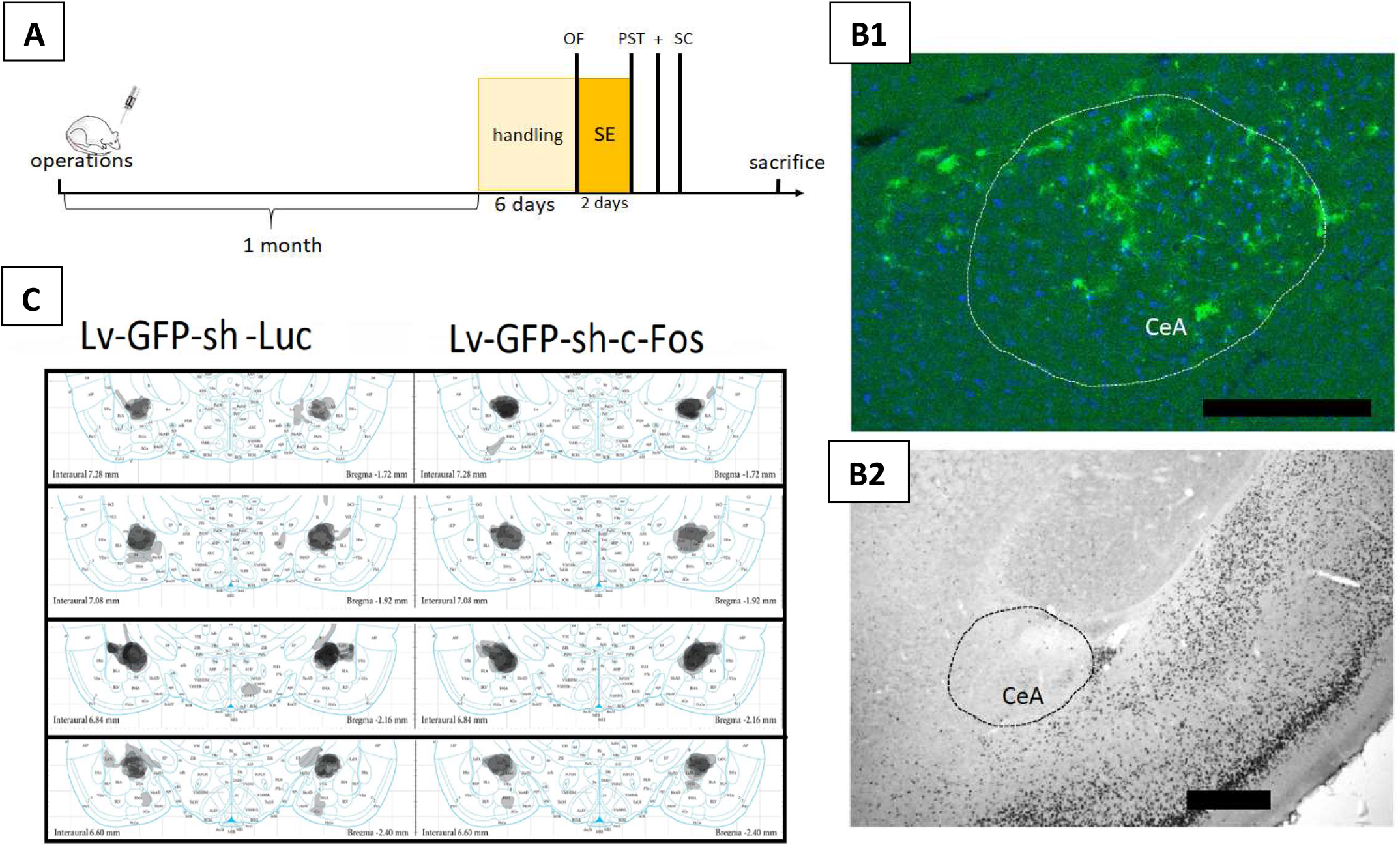
**A** – schematic timeline of experimentation (OF – Open Field Test; SE – Self-Exposure Test; PST – Porsolt Swim Test; + - Elevated Plus Maze Test; SC – sucrose consumption/Anhedonia Test); **B1** – histological representation of LV-sh-c-Fos injection site (GFP–transfected neurons, blue marker – DAPI); **B2** – microphotograph of the same slice as in B1 after c-Fos immunocytochemical screening. For both B1 and B2 microphotographs bar is 500 μm; **C** – Schematic depiction of the injections sites in shLuc and shFos (left and right sides of the panel, consecutively) with an increasing intensity of the grey colouring representing the increasing GFP tagging (graphics adapted from Paxinos & Watson, 2007)

### Behavioural testing

A month after the operations rats were subjected to the behavioural testing (timeline of the experimentation - figure 1A). Firstly for the 6 consecutive days rats were handled by the experimenter. In last day of the handling the Open Field (OF) Test was carried out, with each of the animals for 6 min. The next day the 2 day long Self-Exposure Test (SE) was performed: the first day served as a habituation to the experimental chamber and the second day was an exposure test, when the rats could perform the switch off of the light in the SE chamber for 5 sec, by activating one of 2 photocells inside of the chamber. Ultrasonic vocalisations (USV) were recorded during the two days of the self-exposure test. The apparatus and the setting for SE test were described before (Robakiewicz et al., 2019). A day after the SE test, there was a one 6 min session of Porsolt Swim Test (PST, Porsolt et al., 1979) conducted. There was a one day postponement after PST and the next day Elevated Plus Maze Test (EPM, Pellow et al., 1985; Carobrez and Bertoglio, 2005) was conducted for 6 min for each rat. In the third and the fourth day after the EPM a sucrose (4%) intake test was performed (Anhedonia Test – AT, Strekalova et al., 2004; Frey et al., 2014; Clarkson et al., 2018). More detailed information about the behavioral methods can be found in the supplementary methods section.

## Results

Rats with the blockade of c-Fos have shown no changes in locomortor and explorative behaviors in the OF test. They have covered the same mean and total distances in the field and their mean velocity was similar to the control group (mean and total dist. and veloc., table 1). Frequency in center zone, duration and % of total time spend in that area, was not changed (center zone freq. and durat., table 1). Their anxiety level measured by duration and % of the time spend in the thigmotaxis zone, as well as frequency of entering in it, was not changed comparing to ShLuc group (thigmo. durat. and thigmo. freq., table 1). The only significant difference was in the latency to the first occurrence in the center zone – ShFos rats were faster to enter that zone than the ShLuc rats (center zone laten., table 1 and figure 2). This effect was prominent in the size.

**Figure 2.**
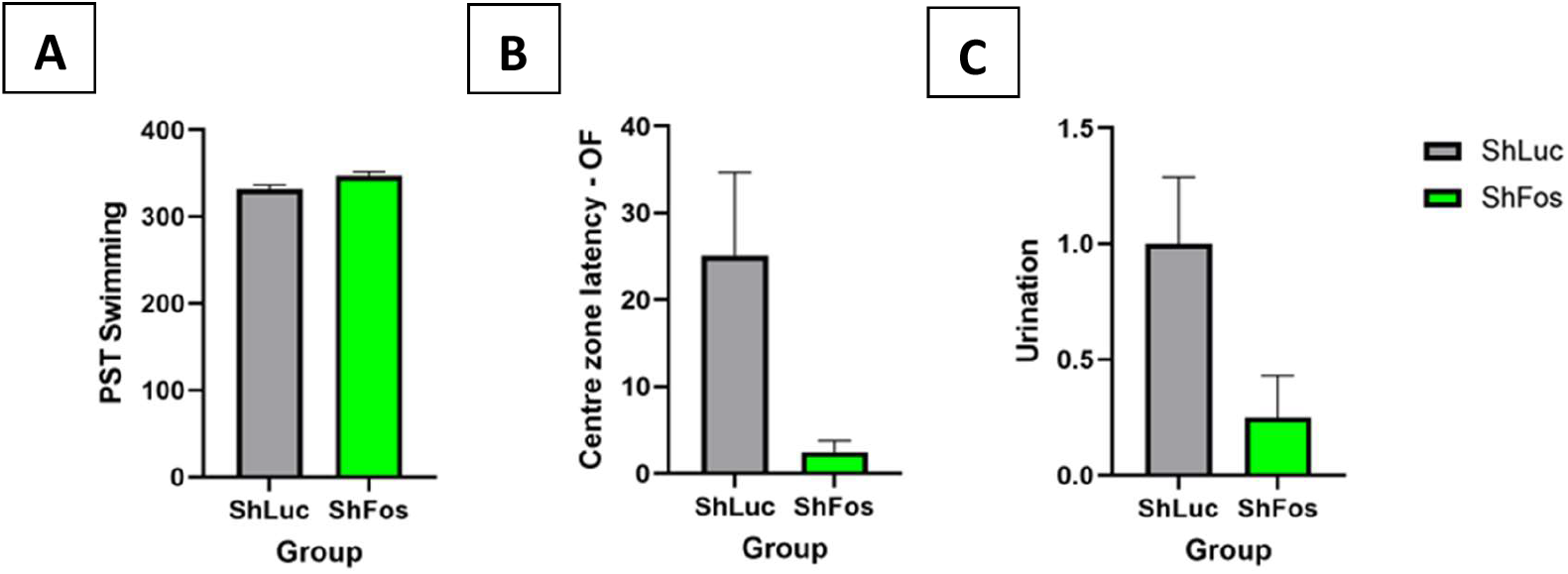
Results of behavioral screening of the operated rats: **A** – Rats from ShFos group were more active in PST; **B** – Rats from ShFos group engaged in more rapid activity in OF then the control group (shorter latencies to the first entrance in the centre zone); **C** – Level of the urination was decreased in ShFos rats comparing to the control group while in EPM test. All group differences are significant on at least *p* < 0.05 level. All graphs represent mean plus the S.E.M.

**Table 1.** *t*-student (*t*) and *U*-Mann-Whitney (*U*) tests comparisons of the behavioral results in shLuc – control group and shFos – experimental group. *M* – mean; *SD* – standard deviation; n – number of animals; *p* – significance level; 95% Cl (LL and CL) – confidence intervals; Cohen’s d – effect size measurement; OF – Open Field Test; SE – Self-Exposure test; PST – Posrolt Swim Test; + - Elevated Plus Maze test; SC – sucrose consumption/Anhedonia Test

|  | ShLuc (n = 9) |  | ShFos (n = 12) |  | *t* | *p* | 95% CI |  | Cohen's *d* |
| --- | --- | --- | --- | --- | --- | --- | --- | --- | --- |
|  | *M* | *SD* | *M* | *SD* |  |  | *LL* | *UL* |  |
| OF dist. mean [cm] | 0.36 | 0.08 | 0.40 | 0.09 | 1.12 | 0.276 | -0.04 | 0.12 | 0.49 |
| OF dist. total [cm] | 3173.37 | 670.82 | 3529.92 | 737.81 | 1.14 | 0.269 | -299.08 | 1012.18 | 0.50 |
| OF mean veloc. [cm/s] | 8.57 | 1.94 | 9.57 | 2.08 | 1.12 | 0.276 | -0.87 | 2.87 | 0.49 |
| OF center zone freq. | 10.89 | 3.86 | 13.08 | 5.79 | 0.98 | 0.338 | -2.48 | 6.87 | 0.43 |
| OF center zone durat. [s] | 12.09 | 6.46 | 16.68 | 13.30 | 0.95 | 0.354 | -5.52 | 14.70 | 0.42 |
| OF center zone [% time] | 3.25 | 1.74 | 4.47 | 3.47 | 0.96 | 0.351 | -1.44 | 3.87 | 0.42 |
| OF thigmo. durat. [s] | 358.38 | 11.58 | 352.14 | 17.92 | -0.91 | 0.375 | -20.61 | 8.13 | 0.40 |
| OF thigmo. durat. [% time] | 96.46 | 1.90 | 94.65 | 3.93 | -1.27 | 0.220 | -4.79 | 1.18 | 0.56 |
| SE hab. 22kHz USV | 10.38 | 7.73 | 14.55 | 11.21 | 0.90 | 0.378 | -5.56 | 13.90 | 0.42 |
| SE test 22kHz USV | 8.78 | 13.38 | 6.92 | 5.52 | -0.44 | 0.667 | -10.76 | 7.04 | 0.19 |
| SE test 50kHz USV | 28.78 | 32.57 | 39.00 | 41.87 | 0.61 | 0.551 | -25.06 | 45.51 | 0.27 |
| SE hab. activity | 21.00 | 11.97 | 15.42 | 8.74 | -1.24 | 0.231 | -15.02 | 3.85 | 0.55 |
| SE test activity | 17.44 | 8.96 | 20.25 | 14.07 | 0.52 | 0.607 | -8.44 | 14.05 | 0.23 |
| PST swimming | 331.74 | 15.46 | 347.23 | 15.19 | 2.29 | **0.033\*** | 1.36 | 29.60 | 1.01 |
| PST immobility | 28.26 | 15.46 | 12.78 | 15.19 | -2.29 | **0.033\*** | -29.60 | -1.36 | 1.01 |
| EPM closed arms | 274.91 | 38.86 | 275.95 | 45.34 | 0.06 | 0.957 | -38.40 | 40.48 | 0.02 |
| EPM open arms | 27.12 | 24.03 | 15.77 | 26.14 | -1.02 | 0.321 | -34.68 | 11.97 | 0.45 |
| EPM urination | 1.00 | 0.87 | 0.25 | 0.62 | -2.32 | **0.032\*** | -1.43 | -0.07 | 1.02 |
| EPM defecation | 1.00 | 1.32 | 0.50 | 1.17 | -0.92 | 0.370 | -1.64 | 0.64 | 0.40 |

**Table 1**
|  | ShLuc (n = 9) |  | ShFos (n = 12) |  | *U* | *p* | 95% CI |  | Cohen's *d* |
| --- | --- | --- | --- | --- | --- | --- | --- | --- | --- |
|  | *M* | *SD* | *M* | *SD* |  |  | *LL* | *UL* |  |
| OF center zone laten. [s] | 25.12 | 28.58 | 2.49 | 4.39 | 13.00 | **0.004\*\*** | -44.66 | -0.60 | 1.20 |
| OF thigmo. freq. | 12.89 | 4.57 | 17.58 | 10.89 | 42.50 | 0.411 | -3.43 | 12.82 | 0.53 |
| SE hab. 50kHz USV | 19.75 | 20.73 | 42.73 | 58.23 | 29.50 | 0.231 | -22.70 | 68.66 | 0.49 |
| SE USV summ. | 64.33 | 38.47 | 98.42 | 93.28 | 40.50 | 0.337 | -35.35 | 103.52 | 0.45 |

ShFos rats have shown unchanged emotional expression level, when it comes to USV emission. They have vocalized the same amount of times in both measured frequencies and in both days of recordings, as well as in the summarized vocalizations from both days and frequencies (hab. and test USV, USV summ., table1). There were a 50 kHz calls and a short 22 kHz calls observed in these experiments, similarly to our previous work with Long-Evans rat strain (Robakiewicz et al., 2019). ShFos rats self-exposure cage activity was on the same level as their ShLuc counterparts (hab. and test activity, table1).

ShFos rats have shown a more active behaviors in PST. Their total time of swimming was longer than the one of the rats from ShLuc group (table 1 and figure 2). Observed effect was prominent in the size.

Anxiety level measured by the time spent in the open arms of EPM was not changed in ShFos rats (table 1). Time spent in closed arms was also similar to the ShLuc group’s time. There was a group difference in urination count – ShLuc rats urinated more often than ShFos rats (table 1 and figure 2). Observed effect was prominent in the size. There were no differences in the defecation level (table 1).

Finally both of the rat groups preferred sweetened solution over tap water – *F* (1, 76) = 104.2, *p* < 0.001. There was a tendency in anhedonia levels, measured by sweetened water consumption - ShFos group consumed less of sweetened water when compared to ShLuc group, but the difference was statistically insignificant – *F* (1, 38) = 3.46, *p* = 0.0704 (ShLuc – M = 88.73, SD = 31.99; ShFos – M = 69.24, SD = 33.55). The effect of the day of experimentation was not significant for sucrose consumption – *F* (1, 38) = 0.001, *p* = 0.975, as well as group and day interaction - *F* (1, 38) = 0.35, *p* = 0.557. The differences were not significant for the group factor - *F* (1, 38) = 0.1, *p* = 0.753, day - *F* (1, 38) = 2.64, *p* = 0.112 and the group and day interaction - *F* (1, 38) = 0.18, *p* = 0.666 for the tap water consumption.

## Discussion

The experiments described in this study are bringing an original input to the knowledge of the role of the c-Fos protein in an emotionally dependent learning in rats. It was proven before, by using the same shRNA construct for the c-Fos blockade, that the protein is involved in a very specific learning and memory processes in auditory cortex (De Hoz et al., 2018) and in the appetitive and aversive learning in the CeA (Lebitko et al., 2021). Most importantly the c-Fos blockade resulted in disruption of the specific experience-dependent memory formation. However it was not known, whether this kind of blockade dependent disruption can be generalised to more rapid types of memory, ones that involve a current stressful stimulation.

ShFos rats fail to show normal level of the stress induced behavioral arrest, prompted by limited escape possibilities and unpleasant water environment in PST (Commons et al., 2017). Similar effects are regularly observed in other experiments with the Wistar rats (Lino-de-Oliveira et al., 2006, Silva et al., 2012), as well as the other rat strains (Lam et al., 2018, Sherwin et al., 2017, Muigg et al., 2007) and in the mice (Yanagida et al., 2016, Nakazawa et al., 2003) and the hamsters (Alò et al., 2014 and 2015) with the usage of an anti-depressive agents and with the possible future therapeutics. Those studies have focused on molecular pathways targeting noradrenaline, serotonin, dopamine, nitric oxide synthase, AMPAergic/GABAergic mechanisms, brain derived neurotrophic factor (BDNF), c-fos mRNA and protein expression, N-methyl-D-aspartic (NMDA) signalling as a sources of an aroused activity levels of the animals in PST. Multitude in those approaches and relatively similar outcome, suggests a generalized function of CeA and particularly c-Fos in it, as a gateway for stress-induced learned behaviors.

Other groups (Regev et al. 2012, Ciocci et al. 2010, Ghiasvand et al. 2011, Yan-Mei et al., 2019, Ventrura-Silva et al., 2020, Poulin et al., 2013, Wiktorowska et al., 2021, Asok et al., 2018, Botta et al., 2015) were aiming at the functional ablation of the CeA and its pathways. In comparison to those studies, here presented approach is less specific and possibly works by blocking the long-term memories formation in all of the transfected neurons, regardless of their neurosecretive and receptive profiles. Although most of the CeA neurons are of an inhibitory GABA-ergic nature, their function is far from homogeneity (Li et al., 2019), making the structure essential for acquisition and expression of the fear based, stress induced learning (Duvarci et. al, 2014, Adraka et al., 2021). This kind of behaviors in PST, were hindered in ShFos animals, possibly indicating on an impaired stress-based learning process.

The basal exploratory and locomortor activity in ShFos rats was intact, as they were showing normal behavior in the OF and in our SE test. Shorter latencies to enter the canter zone of the OF can be interpreted as a sign of low stress levels induced by the novelty (neo-phobia), pointing out again at the disrupted emotional memory formation (Pignatelli & Beyeler, 2019, Botta et al., 2015).

The USV production in ShFos rats was intact by c-Fos blockade. USV generating is realised by the pathways that originate from the brain stem, that are possibly more evolutionary preserved then the Amy itself. The Amy and in particular the CeA is not a main target connection of aversive 22 kHz USV (mostly cholinergic neurons) and appetitive 50 kHz USV (mostly dopaminergic neurons) production pathways (Brudzynski et al., 2021).

Anxiety level measured by the time spend in closed arms of EPM, was not changed in ShFos rats, however the urination level was reduced for them. Micturition is a function of adaptive stress-related activity in the brain, proposed to be regulated by the corticotrophin release factor (CRF) neurons (Chaves et al., 2021, Shimizu et al, 2016). One of the possible ways of ordaining the micturition by emotion-related pathways is from the Berrington’s nucleus that is controlled by the PAG - a main output of the CRF neurons from the CeA (Borelli et al., 2013, Gray and Magnuson, 1992). This CRF-dependent expression of emotions on autonomic level, might be interfered by the ShFos blockade the CeA, in herein presented experiments.

Finally, appetitive, approaching behaviors measured by sucrose consumption are CeA-mediated (Knapska et al., 2013, Lebitko et al., 2021) and seem to be changed in ShFos rats, however the observed difference failed to meet the significance level. The lack of statistically significant effects compared to differences reported by Lebitko and colleagues (2021), might be due to a different species being used in both studies (rats in here vs. mice in theirs) and most probably due to a different experimental design (stronger sucrose solution in theirs – 10% vs. 4% in here, more experimentation days and automated environment in their study, compared to 2 days and human operated testing in here) and due to the methods utilised to measure the behavior (amount of liquid consumed in grams in here vs. licks performed by animal in theirs study).

Altogether our results disclose the c-Fos expression in the CeA to be involved in associative learning, that is based on stress-to-behavior reaction and possibly in the top-down control of micturition from the CeA. The fact that the blockade of c-Fos in CeA on a transcription level, has a significant effect on a behavioral level of rats, constitutes a little piece of information added into a bigger picture of the function of the protein in the affective learning in physiological conditions.

In the future more studies about the CeA-c-Fos activated neurons and their connecting ensembles, possibly in the numerous animal species and with the brain systemic approach that match c-Fos with the other functional agents, could benefit the knowledge of the stress-induced memory more. Surely there are many possibilities of clinical utilisation of such a data.

## Acknowledgements

This study was funded by the National Science Center grant PRELUDIUM/2011/01/N/HS6/04086. I am thankful to the Laboratory of Emotions Neurobiology, the Nencki Institute of Experimental Biology for kindly allowing me to use their microscope.

