## Supplementary methods for "c-Fos protein shRNA blockade in the central amygdala nucleus interfere with rats emotional reactivity on behavioral and autonomic level"

**Supplementary Materials:**

**Supplementary methods**

**Animals:**

Twenty one male Wistar rats, 3 months old, housed in pairs with natural light/dark cycle, with water and food provided *ad libitum* were used. All procedures were approved by the Local Ethics Committee for Animal Experimentation.

**Behavioral testing:**

**OF** arena consisted of a square base measuring 75 x 75 cm, delimited by four walls 75 cm high, all white. Along and across the walls and bottom, there were black lines dividing each of the 5 planes into 16 equal squares. The test was run under reduced lighting conditions to minimise the effects of stress on the results. Each session for a given animal began with inserting the rat into the apparatus, with its mouth facing the side of the wall (always the same one), near the lower right corner of the apparatus. The animals were then allowed to explore the apparatus freely for 6 min. Each of the operated rats participated in this experiment. The animal's behavior was recorded by an HD camera placed above the apparatus and analysed subsequently with EthoVision XT (Noldus Information Technology BV).

For **SE** test, the chambers were controlled and monitored by computer software (PC-Med, Med Associates Inc.). Chambers were 33 x 30 x 27 cm, made of plexiglass (back and front) and the side walls made of aluminium that were equipped with a light bulb (30 lx). Floors were made of metal bars, 0.5 cm in diameter, mounted every 1.7 cm. Each right side aluminium wall had two circular holes, 3 cm in diameter, 14 cm apart, 2 cm above floor level, with photocells to register and count nose-pokes (activity) as radius interruptions. USV were recorded with a high sensitivity condenser microphone (Avisoft Bioacustics), automatically detected and scored on the spectrogram with Avisoft SASLab Pro software (Avisoft Bioacustics). Each chamber was inside of the noise-attenuating wooden box (64 x 38 x 60 cm), equipped with a fan for proper air circulation.

The apparatus for carrying out the forced swim test consisted of a cylinder 100 cm high with a bottom surface closed, 29.5 cm in diameter, made of plexiglass. During the procedure, the apparatus was filled with water with a column height of 39.5 cm at a temperature of 24-26 ° C. Each rat was gently placed on the surface of the water, inside of the **PST** apparatus and remained there for 6 min. Between consecutive experimental sessions, the apparatus was cleaned with an ethanol solution (30%) in order to eliminate odours left after the previous sessions. The animal's behavior was recorded by an HD camera placed in front of the apparatus and subsequently analysed manually by a blind to the animal group experimenter.

The apparatus for carrying out the **EPM** procedure consisted of a cruciform with arms intersecting halfway at an angle of 90°, each 100 cm long. The inner square - the place where the arms were crossed - had dimensions of 10 x 10 cm. The arms in the transverse axis were covered on three sides with walls of 45 cm high. The maze was mounted on 4 legs, 70 cm high, each attached to each of the 4 ends of the arms. Each rat was placed in the center of the apparatus, with its mouth facing the open arm (always the same). The animals were then allowed to explore the apparatus freely for 6 min. The animal's behavior was recorded by an HD camera placed above the apparatus and analysed subsequently with EthoVision XT. The urination and defecation occurrence was counted manually by blind to the experimental group experimenter.

For the Anhedonia Test (**SC**), 2 separate, identical bottles were used for testing each animal, containing separately water and sucrose (4%) in water. Each animal was placed for 24 h in a separate cage with dimensions of 34 x 55 x 19 cm, with food ad libitum available. The rats had access to 2 bottles – one with water and one with sucrose solution (4%) for 24 h. In order to avoid the animal's preference for any of the liquids due to the bottle position, their mutual position was changed after 12 h. Weight of the bottles was measured before the testing and after the testing.
